# Stimulation of the vagus nerve reduces learning in a go/no-go reinforcement learning task

**DOI:** 10.1101/535260

**Authors:** Anne Kühnel, Vanessa Teckentrup, Monja P. Neuser, Quentin J. M. Huys, Caroline Burrasch, Martin Walter, Nils B. Kroemer

## Abstract

When facing decisions to approach rewards or to avoid punishments, we often figuratively go with our gut, and the impact of metabolic states such as hunger on motivation are well documented. However, whether and how vagal feedback signals from the gut influence instrumental actions is unknown. Here, we investigated the effect of non-invasive transcutaneous vagus nerve stimulation (tVNS) vs. sham (randomized cross-over design) on approach and avoidance behavior using an established go/no-go reinforcement learning paradigm (Guitart-Masip et al., 2012) in 39 healthy, participants after an overnight fast. First, mixed-effects logistic regression analysis of choice accuracy showed that tVNS acutely impaired decision-making, *p* = .045. Computational reinforcement learning models identified the cause of this as a reduction in the learning rate through tVNS *(Δα* = −0.092, *p*_*boot*_= .002), particularly after punishment *(Δα*_*Pun*_*=* −0.081, *p*_*boot*_= .012 vs. *Δα*_*Rew*_*=* −0.031, *p* = .22). However, tVNS had no effect on go biases, Pavlovian response biases or response time. Hence, tVNS appeared to influence learning rather than action execution. These results highlight a novel role of vagal afferent input in modulating reinforcement learning by tuning the learning rate according to homeostatic needs.

## Introduction

To survive, organisms must procure energy by approaching options that pay off while avoiding costly options, potentially incurring punishments. Fundamental learning mechanisms have evolved to support this vital optimization of instrumental actions [1–4]. One key challenge is to balance short-term and long-term goals of reward-related behavior. For example, patiently resisting temptation to receive bigger returns later is often beneficial in the longer term. However, in certain bodily states, forfeiting immediate rewards can have negative long-term consequences. For instance, missing out on food in a hungry state [5] can increase the risk of starvation [1], which may profoundly hamper future prospects. Short-sightedness may hence address urgent dietary requirements at the expense of long-term opulence [6]. Despite its obvious evolutionary importance, little is known about how bodily homeostatic needs shape decision making in humans. One plausible candidate for modulatory input onto circuits involved in reward learning would be a caloric feedback signal [7] originating from the gut.

Signals about metabolic and homeostatic state are largely transmitted via the vagus nerve which connects peripheral organs such as the gut with the brain. Vagal afferents terminate in the nucleus tractus solitarii, NTS, [8] a hub further relaying metabolic information to the mid-and forebrain [8,9] including to dopaminergic neurons in the substantia nigra. Along that pathway, vagal afferents have been shown to modulate dopaminergic [10,11], noradrenergic [12], and cholinergic signaling [13]. Accordingly, endogenous stimulation of the gut with nutrients evokes dopamine responses in the dorsal striatum tracking caloric value [14,15]. These dopamine signals are critical for appetitive conditioned learning [10,16,17] as well as motivated behavior [18,19]. Similarly, vagal afferent signaling regulates food intake [16], and stimulation of the vagus nerve has been associated with reduced food intake and decreased weight gain [20]. More broadly, memory function [21], cognitive flexibility [22], and mood [23] are also influenced by vagal signaling. Collectively, these results suggest that vagal signals conveying the metabolic state influence behavior, including reward seeking, via alterations in dopamine.

Until recently, research in humans has been limited by the invasive nature of cervical vagus nerve stimulation (VNS). Nevertheless, VNS has been related to a broad range of behavioral effects such as enhancing cognitive functioning [24] and memory retention [25,26]. VNS also has a role in the treatment of depressive and anxiety symptoms [25]. Lately, non-invasive transcutaneous VNS (tVNS) has become feasible, opening new avenues for research and treatment. It involves transcutaneous stimulation of the auricular branch of the vagus nerve behind the ear. This has been shown to affect projections to the NTS in preclinical studies [27], and studies using tVNS with concurrent fMRI have revealed enhanced activity in the NTS and other interconnected brain regions including the dopaminergic midbrain and nucleus accumbens [28,29]. Moreover, tVNS had similar positive effects on depressive symptoms [30,31], memory retention [32,33], and cognitive performance [34,35] as implanted cervical VNS. Therefore, tVNS provides a promising approach for the investigation of the link between vagal metabolic signals and reward-related decisions.

Previous studies investigating the effects of (t)VNS mainly focused on processes predominantly associated with the noradrenergic system such as cognitive control, fear learning and extinction. Since vagal signals also modulate the dopaminergic system, motivation and reward learning should be shaped by vagal stimulation as well. While response vigor has been linked to dopamine tone [36], potentially reflecting average reward rates, learning via reward prediction errors (RPE) has been linked to phasic dopamine signals [37,38]. The magnitude of RPEs and subsequent learning is also influenced by dopamine tone [39,40]. Perhaps counterintuitively, high dopamine tone reduces the constraint of actions imposed by previous rewards [40] as phasic signals are proportionally smaller (value theory, [41]) or choices rely less on learned values (thrift theory, [2]). Moreover, learning is differentially supported by dopamine depending on the specific task [42], with distinct but interacting circuits underlying learning from rewards or punishment. Consequently, vagal feedback may lead to changes in reward learning, response vigor, and action selection mediated by dopaminergic signaling. Mapping the effects of tVNS onto these motivational facets would shed new light on the endogenous modulation of reward seeking.

In the present crossover study, we applied tVNS (vs. sham) to mimic metabolic signaling via vagal afferents and tested its effects on reward learning, which may be mediated by changes in dopamine levels. Reward learning was probed with a valence-dependent go/no-go learning paradigm established by Guitart-Masip et al. [42] delineating instrumental action learning and Pavlovian control. In line with the value and thrift theories of dopamine and our recent work corroborating these predictions [40], we hypothesized that during tVNS participants’ performance would be impaired, as increased dopamine tone reduces the impact of phasic RPEs. We then used computational reward-learning models to investigate specific valence-or action-dependent changes in performance. In a previous study, Guitart-Masip et al. [43] reported a decrease in Pavlovian bias by increased dopamine levels (after L-DOPA administration) thereby improving performance in incongruent action-valence combinations while reducing performance in congruent action-valence combinations. Hence, we expected tVNS to lead to a similar attenuation of the Pavlovian influence on instrumental learning. In addition, we explored potential effects of tVNS on greater go response rates or faster response time, which would be indicative of heightened vigor.

## Methods

### Participants

In total, 44 individuals participated in the study. They were physically and mentally healthy, German speaking, and right-handed, as determined by a telephone interview. For the current analysis, five participants had to be excluded (n=4: did not complete both task sessions, n=1: did not make any go response). Thus, we included 39 participants (23 female, *M*_*age*_= 25.5 ±4.0 years; *M*_*BMI*_= 23.0 ±3.0 kg/m^2^). The institutional review boards of the University of Tübingen approved the study and we obtained informed consent from all participants prior to taking part in the experiment.

### Experimental procedure

Participants were required to fast overnight (i.e., >8h) before both experimental sessions. Sessions were conducted in a randomized, single-blind manner as the experimenter was not blind to the stimulation condition (for information on the device, see SI). Nevertheless, participants were close to chance in guessing the correct condition (60%; *p*_binomial_= .049) suggesting that blinding was effective. Sessions started between 7.00am and 11:00am and lasted about 2.5h in each. After participants arrived for the first session, they provided written informed consent. Next, we collected anthropometric and state-related information (see SI) before the tVNS electrode was placed on the left ear targeting the auricular branch of the vagus nerve. In line with the stimulation procedure by Frangos et al. [28], the electrode was located at the left cymba concha for tVNS and at the left earlobe for sham stimulation. Stripes of surgical tape served to secure the electrode in place. We determined individual stimulation strength for every session separately using pain VAS ratings (“How intensely do you feel pain induced by the stimulation?” ranging from 0 (“no sensation”) to 10 (“strongest sensation imaginable”). Stimulation was initiated at an amplitude of 0.1mA and increased by the experimenter in 0.1-0.2mA steps. Participants rated the sensation after every increment until ratings settled around 5 corresponding to “mild prickling”. Then, the stimulation continued throughout the task block according to the stimulation protocol of the device (i.e., alternating blocks of stimulation on and off for 30s each). Within this block, participants completed a food-cue reactivity task (∼20min) and an effort allocation task (∼40min) before the learning task.

After completing state-related questions, participants received rewards according to task performance and compensation (either as 32€ or partial course credit). Both visits followed the same standardized protocol.

### Paradigm

We hypothesized that tVNS affects reinforcement learning via changes in dopaminergic neurotransmission. However, increases in dopaminergic transmission are not universally translated into increases in performance as expected changes critically depend on the task. Due to the well-known characteristics of the dopaminergic circuit [44,45], we sought to disentangle effects of tVNS on action-or valence-dependent learning. To this end, we used a previously established go/no-go reinforcement learning task [42,43,46]. In this task, participants learn state-action contingencies and receive rewards or punishments. Each trial consisted of three stages (Figure 1). First, participants saw a fractal cue (state) out of a set of four different fractals per session. These fractals were initially randomized to one of the four possible combinations of the go × win two-factorial design of the task. Second, participants had to complete a target detection task and either respond by pressing a button (go) or withhold their response (no-go). Third, they saw the outcome of the state-action combination, which was either a win (5 cents), punishment (−5 cents), or an omission (no win/punishment, 0 cents). Using trial and error, participants had to learn which action following each fractal was best in terms of maximizing wins or minimizing losses.

**Figure 1:**
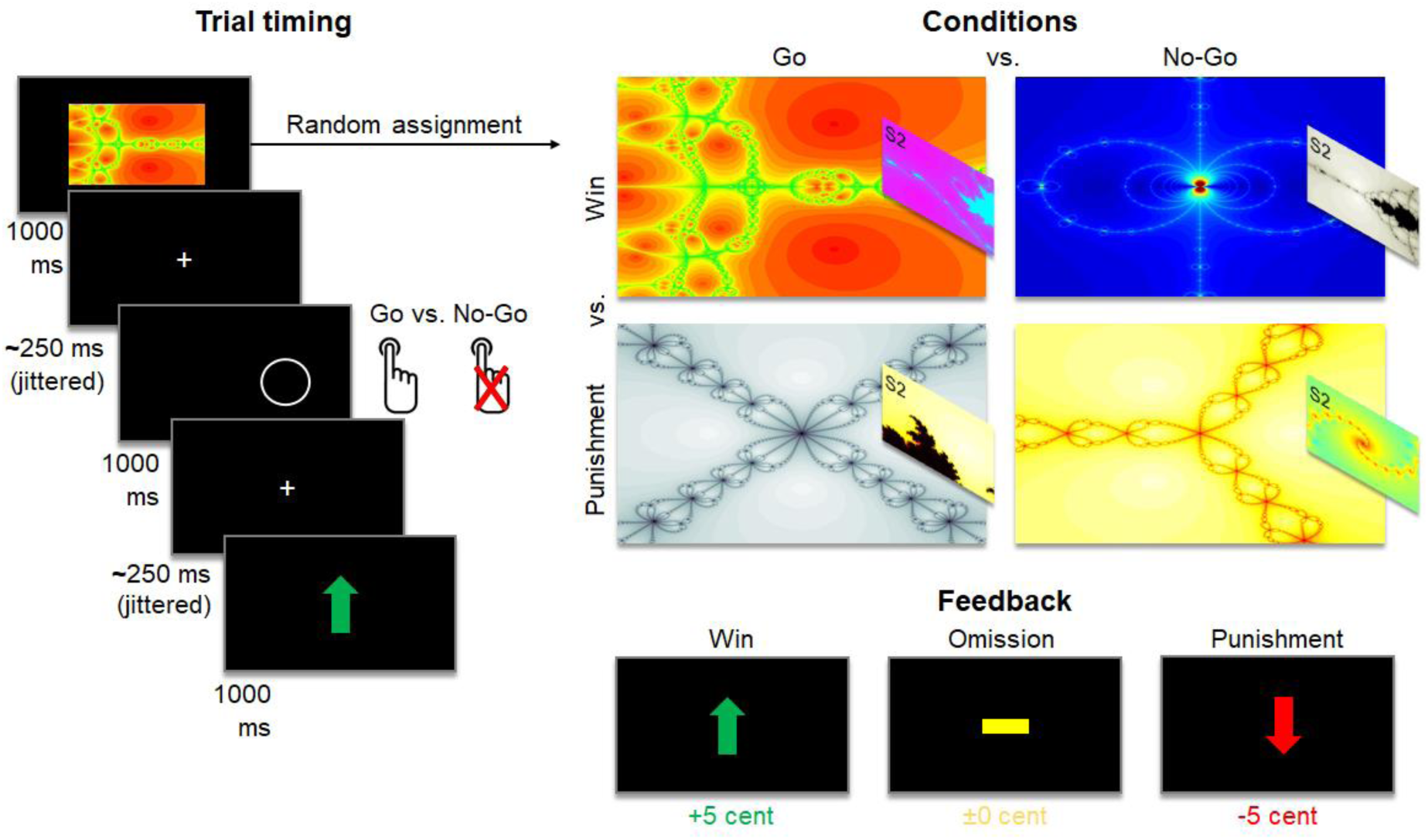
Schematic summary of the go/no-go reinforcement learning task. To maximize the total payoff, participants had to learn which action (go vs. no-go) during the target detection stage following a given fractal was associated with the best possible outcome (i.e., receiving reward or avoiding impending punishment). These contingencies were randomly assigned and had to be learned by trial and error. Insets illustrate that different fractals were used in Session 2 (S2).

**Figure 2:**
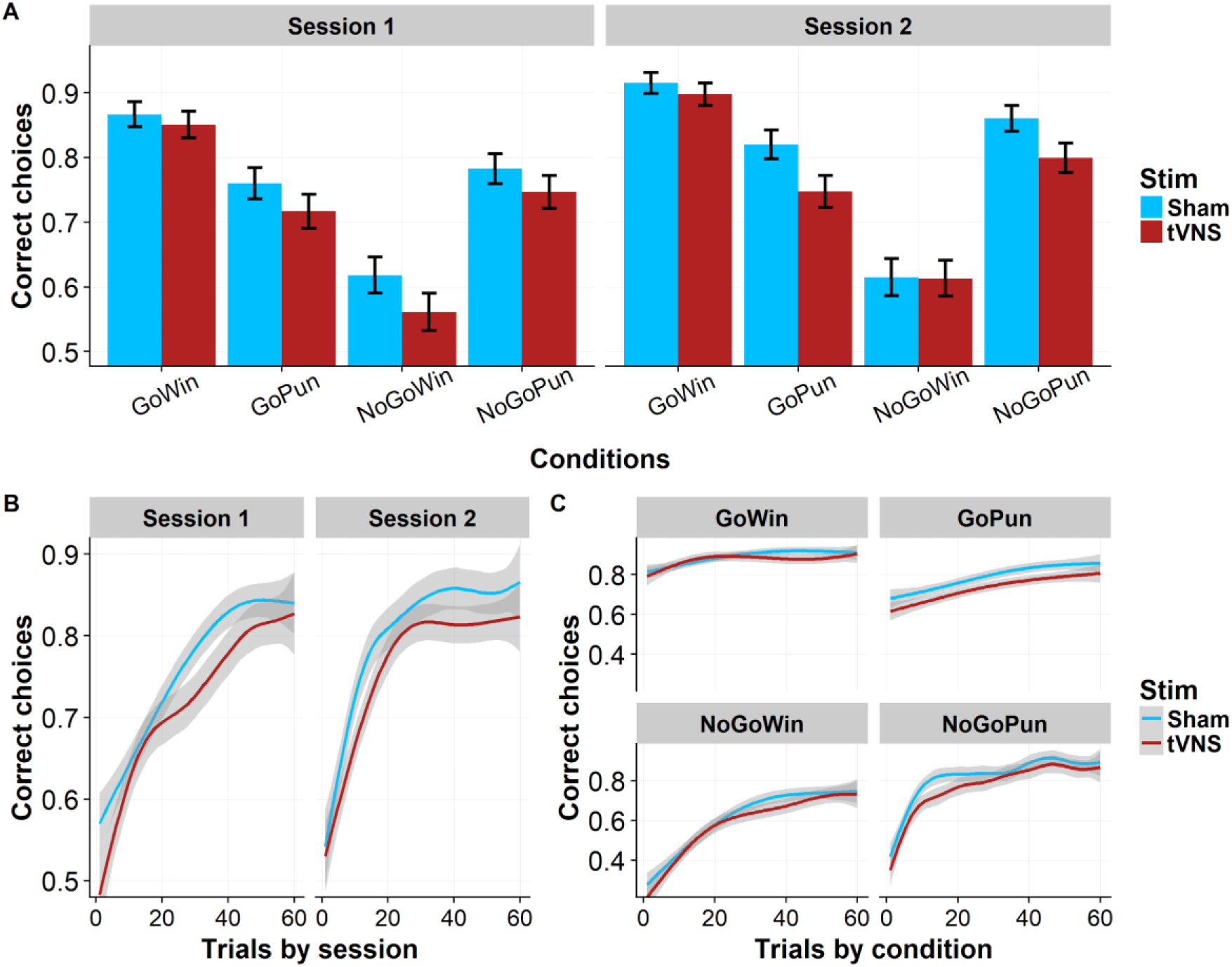
Choice accuracy is reduced in the tVNS condition compared to sham stimulation. A: Mean choice accuracy for tVNS and sham stimulation in each session and condition. Error bars depict 95% confidence intervals. B: Choice accuracy for tVNS and sham stimulation over trials separated by session indicate stronger tVNS-induced reduction of choice accuracy in session 1. C: Choice accuracy for tVNS and sham stimulation over trials separated by condition do not suggest action-or valence-specific effects of tVNS.

Outcomes were presented probabilistically. Thus, participants had 80% chances to win after correct state–action sequences, 20% chances to win after incorrect sequences for rewarded trials as well as 80% chances to avoid losses after correct, and 20% chances to avoid losses after incorrect sequences for punished trials. Participants were instructed about the probabilistic nature of the task and that either go or no-go responses could be correct for a given fractal. There was no change in the contingencies over time. To ensure that participants understood the task, they were queried before starting the task. In total, the task included 240 trials (60 per condition) and took 15min to be completed.

## Data analysis

### Full mixed-effects analysis of the go/no-go reinforcement learning task

To estimate the effects of tVNS on choice accuracy (go vs. no-go), we defined a full mixed-effects analysis as implemented in HLM [47]. Effects of the conditions were modeled by predicting if a given choice (Bernoulli distribution) was correct based on the regressors go (dummy coded), win (dummy coded), and the interaction term go × win in a generalized linear model. To assess tVNS effects, the model included terms for the stimulation condition (dummy coded, 0=sham, 1=tVNS) and interactions of the stimulation term with the condition regressors (i.e., stimulation × win, stimulation × go, stimulation × go × win). Furthermore, we included a log-transformed trial regressor capturing improvements in accuracy across trials. At the participant level, we calculated two models that included random effects for all intercepts and slopes: model 1 controlled only for order, whereas model 2 additionally controlled for gender and BMI. We also tested an additional interaction term stimulation × trial but found that the coefficient estimate was highly correlated with the stimulation main effect. Thus, we excluded this term to avoid redundancy. All other random effects were complementary and showed significant between-subject variance (*p*<.001). Analogous to using expectation maximization (EM) in the computational model, we obtained empirical Bayes estimates, which take group-level distributions into account, as individual estimates of tVNS effects.

### Reinforcement learning model

To dissociate which facet of instrumental action learning was altered by tVNS, we fit reinforcement-learning models to participant’s behavior starting with the winning model detailed in Guitart-Masip et al. [42,43] as standard model (for details and equations, see SI). Models were fit using hierarchical EM as described by Huys et al. [48]. Model fit was assessed using group-level integrated BIC (iBIC, [48]) where model fit and model complexity across all measurements are taken into account. As better group-level fit may be driven by large improvements in few participants, we additionally used likelihood-ratio tests to determine the best fitting model for each session. To ensure stability of individual estimates, we used 10 EM initializations to calculate the mean and the coefficient of variation. Furthermore, we assessed recovery of observed behavior based on simulations with estimated parameters.

### Statistical analysis and software

We assessed all tVNS effects using a significance threshold of *p* < .05 (two-tailed) and corrected for multiple comparisons across the five parameters in the computational model analysis using Bonferroni correction. We also planned correction across condition-specific interaction terms in the mixed-effects model, but they did not reach uncorrected significance. To account for non-normal distributions of parameters from the computational model, differences in parameter estimates between the tVNS and sham condition were tested using bootstrapping (1000 resamples). We performed data analyses with Matlab v2016a (computational model) or HLM v7 (mixed-effects models) and data visualization with R v5.0.1 and Deducer [49].

## Results

### tVNS reduces choice accuracy across conditions

We first analyzed the performance of participants by estimating effects of reward valence, required action, and stimulation on accuracy in a full mixed-effects model. In line with previous studies, accuracy was higher in conditions requiring a go response (*t* = 5.93, *p* < .001), whereas reward valence only influenced accuracy in interaction with the required action (valence: *t* = 0.83, *p* = .412, valence × action: *t* = 7.198, *p* <.001). In other words, participants performed worse in the *go-punishment* and *no-go-win* conditions in which Pavlovian biases (*approach reward*; *avoid punishment*) and instrumental behavior were incongruent.

Next, we assessed main and interaction effects of tVNS vs. sham stimulation on choice accuracy. Across conditions, tVNS reduced accuracy (*t* = −2.08, *p* = .045; model uncorrected for BMI and sex: *t* = −1.98, *p* = .055). However, we observed no interaction effects with action (*t* = −0.46, *p* = .646) or valence (*t* = 0.78, *p* = .44). As task-dependent improvement across sessions is common in reinforcement learning tasks, we controlled for stimulation order (sham/tVNS first) in the analyses. Order of stimulation modulated stimulation slopes (*t* = −3.60, *p* < .001) with stronger impairments in overall performance if tVNS was applied first. Notably, acute tVNS-induced reductions in performance did not lead to deficits in the second session with higher day-to-day improvements in the group that received tVNS first (*t* = 2.05, *p* = .048).

### tVNS reduces the learning rate in a computational model of behavior

To further characterize which learning processes were affected by tVNS leading to impaired performance, we fitted a computational reward-learning model [42] using an EM algorithm to empirically regularize parameter estimates. We estimated five parameters controlling choices over time: learning rate, reward sensitivity, go bias, Pavlovian bias, and noisiness of choices for each session and calculated differences between tVNS and sham sessions. Simulated data based on individually estimated parameters corresponded well with observed data (Figure 3C-E) and parameter estimates were sufficiently stable across the 10 reruns with median coefficients of variation between .002 and .030 for the five parameters.

**Figure 3:**
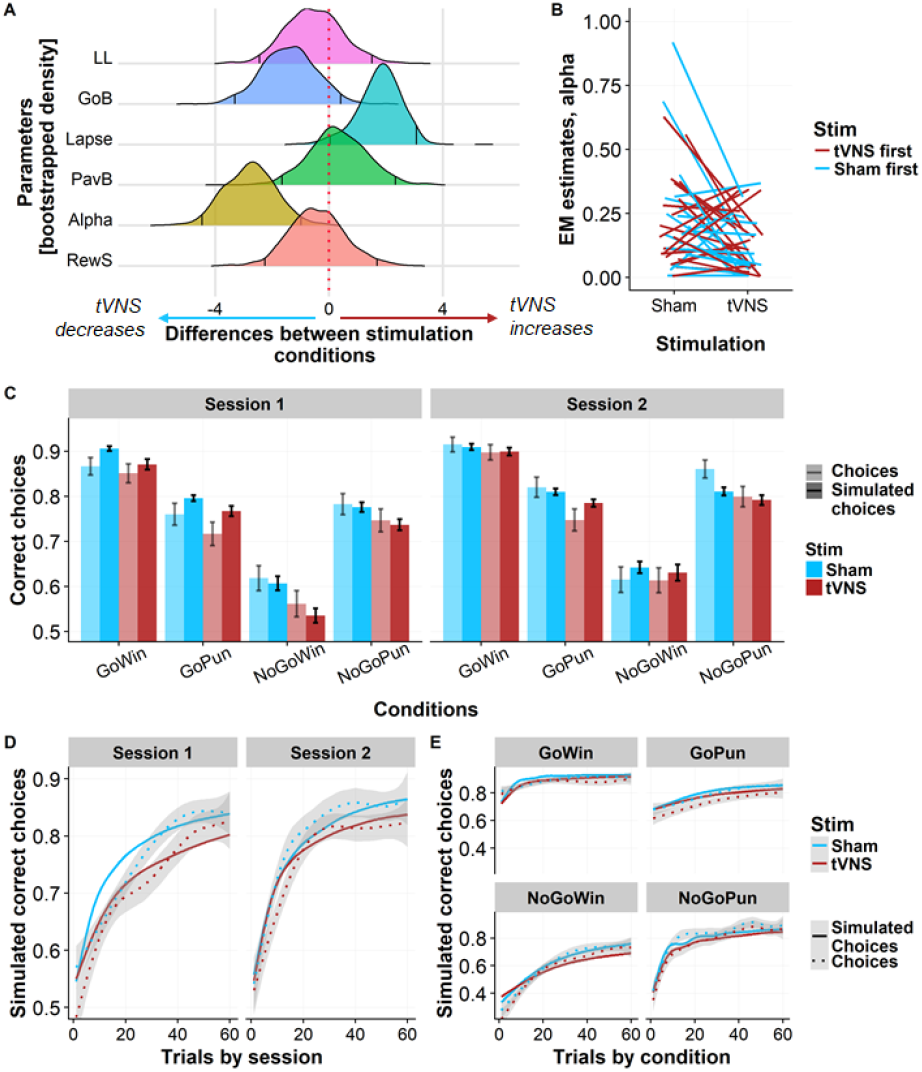
Reduced choice accuracy in the tVNS condition is driven by a reduced learning rate and increased choice stochasticity in the 5-parameter computational model. A: Bootstrapped density plots of the differences in individual parameter estimates between tVNS and sham stimulation. Lines indicate 95% confidence intervals. B: Individual changes in the learning rate indicate mainly a reduction of high baseline learning rates after tVNS. Stimulation in session 1: blue = sham, red = tVNS C: Mean choice accuracy simulated from individual parameter estimates recover stimulation effects in each session and condition. Error bars depict 95% confidence intervals. Transparent bars depict choices, dark bars simulated choices. D: Choices simulated from individual parameter estimates recover participants’ choice patterns and stimulation effects for sessions. E: Recovered choice patterns indicate no difference in stimulation effects depending on valence or action. LL = Log-Likelihood, GoB = Go bias, PavB = Pavlovian bias (π), RewS = Reward sensitivity (ρ)

Impaired performance during tVNS was mainly reflected in a reduced learning rate alpha (Figure 3A; Δα = −0.092, *p* = .009, *p*_*boot*_= .002, corrected for stimulation order: *t* = −2.741 *p* = .009; α_Bonferroni_= .01). Additionally, participant’s choices in the tVNS condition were ‘noisier’ and less dependent on learned action values (Δξ = 0.035, *p* =.086, *p*_*boot*_= .050), although only nominally significant before correction for multiple testing.

Valence-specific effects of tVNS may be captured by modeling separate parameters for rewards and punishments. Therefore, we also built an extended 6-parameter model assuming separate learning rates. While the 6-parameter model provided a more parsimonious account at the group level (ΔiBIC = 263), it only improved individual model fits for 27 out of 78 sessions. In line with the 5-parameter model, tVNS reduced learning rates. Interestingly, this effect was primarily driven by a decrease in the punishment condition (see SI).

We also explored more complex models by additionally separating reward sensitivity and/or Pavlovian bias for reward and punishment as previously described [46]. However, these models did not provide a more parsimonious account at the individual level compared to the simpler models and individual estimates became increasingly unstable across iterations precluding their use to reliably estimate within-subject stimulation effects (see SI).

### tVNS effects are associated with body weight

Behavioral effects of dopamine increases are known to depend on baseline dopamine tone. One well-established correlate is BMI [50,51] and tVNS effects on general accuracy (*t* = 1.987, *p* = .055), as well as on learning rate (*t* = 2.351, *p* = .024) depended partly on participants’ BMI. More specifically, tVNS reduced the speed of acquisition more strongly in participants with a low (healthy) BMI, who have a higher dopamine tone in the striatum compared to overweight participants.

### tVNS does not affect response time

Lastly, we estimated effects of tVNS on response time as an indicator of alterations in response vigor. However, no significant changes in response time were observed (*t* = 0.826, *p* = .414, Figure S.2). This further corroborates that impaired performance was mediated by slowed learning and not by altered action selection.

## Discussion

The vagus nerve rapidly transmits interoceptive signals to the brain. It thereby confers information such as metabolic state and modulates the dopamine system. Here, we investigated changes in instrumental reinforcement learning, which is critically dependent on dopamine, after emulating vagal feedback signals using tVNS. Importantly, we found that tVNS reduced overall accuracy of choices driven by a slowed acquisition of action contingencies, predominantly for punishments. In contrast, action-or valence-specific biases were unaffected by tVNS. In line with the value hypothesis of dopamine [40,41], the observed attenuation of learning rates may be explained by an increase in dopamine tone leading to a lower signal-to-noise ratio of phasic dopamine signals [40,41]. Thus, using the novel non-invasive stimulation of the vagus nerve, our results provide evidence that metabolic feedback signals may alter reward learning by tuning the speed of acquisition according to homeostatic needs.

Vagal feedback signals evoked by tVNS acutely impaired choice accuracy and reduced learning rates in valenced go-/no-go learning. This is in agreement with the value theory of dopamine and with studies showing that the impact of phasic RPE signals on actions depends on dopamine tone [40,41]. In short, increased dopamine tone leads to a comparably smaller signal-to-noise ratio if phasic signals are unaffected and to reduced action control via phasic outcome signals. Accordingly, reduced learning after L-DOPA administration has been reported in patients with Parkinson’s disease [52,53] as well as healthy participants [54]. Similarly, reduced learning rates are in line with the dopamine overdose hypothesis [55], as tVNS reduced learning rates more strongly in participants with a healthy weight, who were previously shown to have a higher dopamine tone in the striatum compared to overweight and obese individuals [50,51]. This may indicate that tVNS-induced increases in dopamine tone led to reduced learning rates predominantly in participants with optimal baseline dopamine function.

In addition to slower learning, impaired performance in the task may also be caused by increased choice stochasticity as predicted by the thrift hypothesis of dopamine [2]. Here, increased dopamine tone would indicate heightened average reward and energy availability [36], consequently leading to more exploration reflected in an uncoupling of learned value and choice [56]. In agreement with the thrift theory, we found that tVNS was associated with increased decision noise [57,58]. However, the increase was not significant after correction for multiple testing and not consistent across models suggesting limited effects at best. These discrepancies may partly be explained by different parameterizations of the computational models. Whereas choice stochasticity is often captured with the temperature parameter in the softmax function, this parameter is separated in the common task model proposed by Guitart-Masip et al. to differentiate reward sensitivity from actual decision noise. Comparably, Guitart Masip et al. [43] did not report changes in the noise parameter in the same task using the same parametrization after L-DOPA administration. Collectively, these results suggest that tVNS primarily affects action–contingency learning and not solely noise in value-based decisions.

In contrast to our hypothesis, tVNS did neither affect response-specific biases such as Pavlovian or go biases nor response times in any condition. In previous studies, pharmacologically-induced increases in tonic dopamine modulated Pavlovian [43] or motivational biases [59] and differentially affected learning from rewards versus punishments [60,61]. Although, we observed that punishment learning, but not reward learning was significantly reduced, there was no significant interaction between valence-dependent learning rates and tVNS effects. Thus, tVNS-induced effects were generally independent of valence or the required action, which is not in line with previous pharmacological interventions increasing dopaminergic transmission. One possible explanation is that tVNS affects multiple transmitter systems and their interplay might lead to mixed behavioral alterations. However, many dopaminergic drugs such as L-DOPA also act on other transmission systems [62] suggesting that this is an insufficient explanation. Another possibility is that modulatory effects of tVNS are more confined within the motivational circuit compared to systemic drug administration. For example, it is conceivable that tVNS could alter the balance between fast reinforcement learning, primarily linked to the amygdala, and slow reinforcement learning, primarily linked to the striatum. It has been shown that chronic tVNS increases functional connectivity between the amygdala and the prefrontal cortex in depressed patients [31] whereas VNS acutely reduces amygdala-evoked responses in the prefrontal cortex of rats [63]. Thus, future research may help to resolve these questions by detailing corresponding alterations in motivational circuits as our study leads to testable predictions about shifting the balance more towards slow striatal reinforcement learning [64]. To conclude, tVNS appears to reduce the speed of learning without altering action-related processing, but more research is needed to establish differences in these processes between endogenous versus exogenous modulations of dopaminergic transmission.

While behavioral effects of tVNS, including reduced accuracy and learning rates, can be explained by modulations of dopamine, our study has limitations. First, although we focused on dopamine, tVNS is not specifically targeting only one neurotransmitter. tVNS has also been associated with heightened noradrenergic signaling via the locus coeruleus leading to improved memory performance by increasing arousal and attention [33]. Nonetheless, phasic noradrenaline signals have also been shown to track unsigned prediction errors (“surprise”) [65,66]. Surprise signals are critical for learning and, accordingly, treatment with an noradrenaline reuptake inhibitor was associated with comparable baseline-dependent changes in learning rates [67]. Since action contingencies are fixed throughout the task, it is not possible to dissociate dopaminergic and noradrenergic processes acting via signed (reward) or unsigned (surprise) prediction errors, respectively. Moreover, as dopamine is the precursor of noradrenaline, future studies disentangling both systems are necessary. Second, the within-subject cross-over design offers increased statistical power to detect stimulation effects, especially considering baseline dependence. Nonetheless, repeated completion of the task may have affected performance and modulated tVNS effects. We accounted for order effects in the statistical analyses, but replication in independent groups would be preferable.

To summarize, we showed that vagal signals impair choice accuracy by acutely reducing learning speed in a reinforcement learning task. These findings could indicate that vagal afferents modulate dopamine tone and, in accordance with the value theory of dopamine, slower acquisition may be due to a reduced signal-to-noise ratio of evoked phasic dopamine. We conclude that how much we learn from rewards and punishments may depend on the metabolic state signaled by the vagus nerve. Thereby, rapid learning which actions in each state lead to future reward or punishment could be facilitated during a hungry state compared to a less deprived state. Critically, this behavioral flexibility with respect to the current metabolic state was less pronounced in overweight participants with a lower baseline dopamine tone, which is in line with the reported reduced sensitivity to peripheral metabolic feedback [68]. Furthermore, reported anti-depressant effects of tVNS may partly rely on reduced learning, especially from punishments, as this may compensate for the reported increased punishment sensitivity in depressed patients [46,69]. More broadly, reduced dependence on learned contingencies may also offer the possibility to prevent over-reliance on learned action-outcome combinations and encourage exploration. In turn, this could lead to greater behavioral flexibility that may be advantageous in many naturalistic environments.

## Acknowledgement

We thank Anni Richter for sharing the German task instructions with us. The study was supported by the University of Tübingen, Faculty of Medicine fortune grant #2453-0-0. VT, CB, & NBK received salary support from the University of Tübingen, Faculty of Medicine fortune grant #2453-0-0. MPN received salary support from the Else Kröner-Fresenius Stiftung, grant 2017_A67. QH is supported by the UCL NIHR BRC.

## Author contributions

NBK was responsible for the study concept and design. VT implemented the task. CB & MPN collected data under supervision by MW & NBK. AK, QJMH, & NBK conceived the method including statistical and computational models. AK & NBK processed the data and performed the data analysis. AK & NBK wrote the manuscript. All authors contributed to the interpretation of findings, provided critical revision of the manuscript for important intellectual content and approved the final version for publication.

