## Supporting Information for "Stimulation of the vagus nerve reduces learning in a go/no-go reinforcement learning task"

Calwerstr. 14, 72076 Tübingen, Germany

### Experimental procedure

After obtaining informed consent, we measured participant's pulse, weight and height as well as waist and hip circumference according to WHO guidelines [1]. Moreover, participants reported their last meal and drink and female participants also reported oral contraceptive use and the beginning of their last menstrual cycle. They also indicated which breakfast they would like to receive after completing the study as part of another motivational task. Using the joystick on an XBox-360 controller (Microsoft Corporation, Redmond, WA), participants responded to state questions presented on a computer that were scored using visual analogue scales (VAS). Mood state items were derived from the Positive and Negative Affect Scales (PANAS; Watson, Clark, & Tellegen, 1988) and complemented by questions on metabolic state. These state VAS ratings were completed at three timepoints (t0 = before stimulation and the task block, t1 = after the task block, yet before breakfast, t2 = after breakfast).

By the end of the task block, participants completed a second set of state VAS. Then, the tVNS device was removed and participants received a snack and their breakfast. For breakfast, a 10-min break was scheduled, but most participants finished eating before the end of the allotted break. A last set of state VAS ratings and a brief questionnaire about physical activity during the past 7 days were answered before participants received their remuneration.

### tVNS device

To stimulate the auricular branch of the vagus nerve, we used the *NEMOS*® stimulation device (cerbomed GmbH, Erlangen, Germany). These devices have been previously employed in clinical trials [2,3] and proof-of-principle neuroimaging studies [4]. The stimulation protocol for the *NEMOS*® is preset to a biphasic impulse frequency of 25 Hz with a stimulation duration of 30 s, followed by a 30 s stimulation pause. The electrical current is transmitted by a titanium electrode placed at the cymba conchae (tVNS) or earlobe (sham) of the left ear [4]. To match the subjective experience of the stimulation, intensity was determined for each participant and each condition individually to correspond to mild pricking (tVNS: $M_{tVNS}$ = 1.21 ±0.43; 0.2-1.9 mA; sham: $M_{sham}$ = 1.92 ±0.67; 0.5-3.1 mA).

**Reinforcement learning model**

Participants learn stimulus (*s*) specific action (*a*) values (*Q*) that are updated at each trial *t* according to the Rescorla-Wagner rule as follows:

$$Q_t(s_t, a_t) \, = \, Q_{t-1}(s_t, a_t) + \alpha(\rho r_t \, - \, Q_{t-1}(s_t, a_t)),$$

with learning rate alpha (α ϵ [0,1]), reward sensitivity rho, a positive free parameter quantifying the individual importance of reward and obtained rewards $r_t$ coded as -1 in case of punishment, 1 in case of reward and 0 if participants received neither reward nor punishment. Further, agents learn action-independent values (V) of each state updated after the same rule indicating if a stimulus is associated with punishments or rewards.

$$V(s_t) \, = \, V_{t-1}(s_t) + \alpha(\rho r_t \, - \, V_{t-1}(s_t)),$$

Action values (Q) and stimulus values (V) at each trial are used to compute action weights as follows:

$$W_t(a,s) = \begin{cases} Q_t(a,s) + b + \pi V_t(s), & a = go \\ Q_t(a,s), & else \end{cases}$$

Where $b$ is a free parameter that reflects a constant bias to choose the go option. The influence of Pavlovian tendencies (e.g. increased go behavior in potentially rewarding situations and avoidance in aversive situations) is parameterized by $\pi$, a positive free parameter. The Pavlovian parameter inhibits the go tendency in conditions that are associated with punishments and thus have negative learned state-values (V), while it increases go tendencies in conditions associated with reward and positive state-values. Consequently, this leads to impaired learning in incongruent (e.g. go-to-avoid punishment) trials.

The action at each trial is selected based on action probabilities that are estimated by passing action weights $(W)$ through a softmax function and adding a noise parameter $(lapse, \xi \in [0,1])$ modulating the influence of learned expectations on subsequent decisions.

$$p(a_t|s_t) = \left[\frac{\exp\big(W(a_t|s_t)\big)}{\sum_a \exp(W(a'|s_t]}\right](1-\xi) + \frac{\xi}{2}$$

Subsequently, we fit three further models to disentangle possible effects depending on reward valence by estimating either learning rate, learning rate and reward sensitivity, or learning rate, reward sensitivity and pavlovian bias for reward and punishment conditions separately.

To fit models with expectation maximization, individual parameters as well as the underlying group distribution parameters are estimated iteratively. The current group distributions were used as priors to estimate individual level parameters using Laplace approximation in the E-step. Consequently, in the M-step, group-level distributions were updated based on the new individual parameter estimates and their uncertainty. Repeated sessions were treated as independent measurements and one underlying distribution was fit over all participants and measurements. Reward sensitivity and Pavlovian bias parameters were log transformed and learning rate and noise parameters were transformed using the inverse sigmoid function to ensure theoretical parameter constraints.

**Computational model with six free parameters**

The 6-parameter model provided a better model fit on the group level (ΔiBIC = 263), although on an individual level, model fit was only significantly improved for 27 out of 78 runs. Nonetheless, stability of individual parameter estimates was sufficient (median coefficient of variation in the range between 0.004 - 0.074 for the six parameters). Subsequent estimation of tVNS effects revealed that the slower learning rate during tVNS stimulation was predominantly driven by a decrease of the learning rate in the punishment condition ($\Delta\alpha_{pun}$ = -0.081, $p$ = .019 , $p_{boot}$ = .012, corrected for order: $t$ = -2.516, $p$ = .016) while decreases of alpha in reward conditions were less pronounced and non-significant ($\Delta\alpha_{rew}$ = -0.031, $p$ = .219 $p_{boot}$ = .21, corrected for order: $t$ = -1.244, $p$ = .211). However, the interaction between stimulation × valence for the learning rate was not significant, $F(1,37) = 1.975$, $p$ = .168, indicating only weak specificity of the tVNS effect on punishment learning. In contrast to the 5-parameter model, tVNS did not affect choice stochasticity in the extended model (Δξ = -0.0031, $p$ =

.863, $p_{boot}$ = .941). Again, stimulation effects on performance were recovered in the averaged simulated data (Figure S.1. C-D).

### Computational model with more than six free parameters

Model comparisons revealed the most complex model (the parameters learning rate, reward sensitivity, and pavlovian bias were separated for reward and punishment conditions) to be the most parsimonious (ΔiBIC = 320, compared to the 5-parameter model, ΔiBIC = 57, compared to the 6-parameter model), but model fit was only significantly improved for 24 out of 78 runs (31%) compared to the 5-parameter model. Moreover, individual parameter estimates across EM initialization using different random seeds were not stable preventing analysis of tVNS effects. Separating only one more parameter (reward sensitivity or pavlovian bias) in addition to the learning rate did not further improve model fit markedly ($\Delta iBIC_{RS}$ = 283, $\Delta iBIC_{PavB}$ = 215, both compared to the 5-parameter model). Simulations of all further models were comparable indicating that additional parameters did not explain additional patterns in the data. Consequently, we did not estimate tVNS effects for any of the more complex models.

### Figures

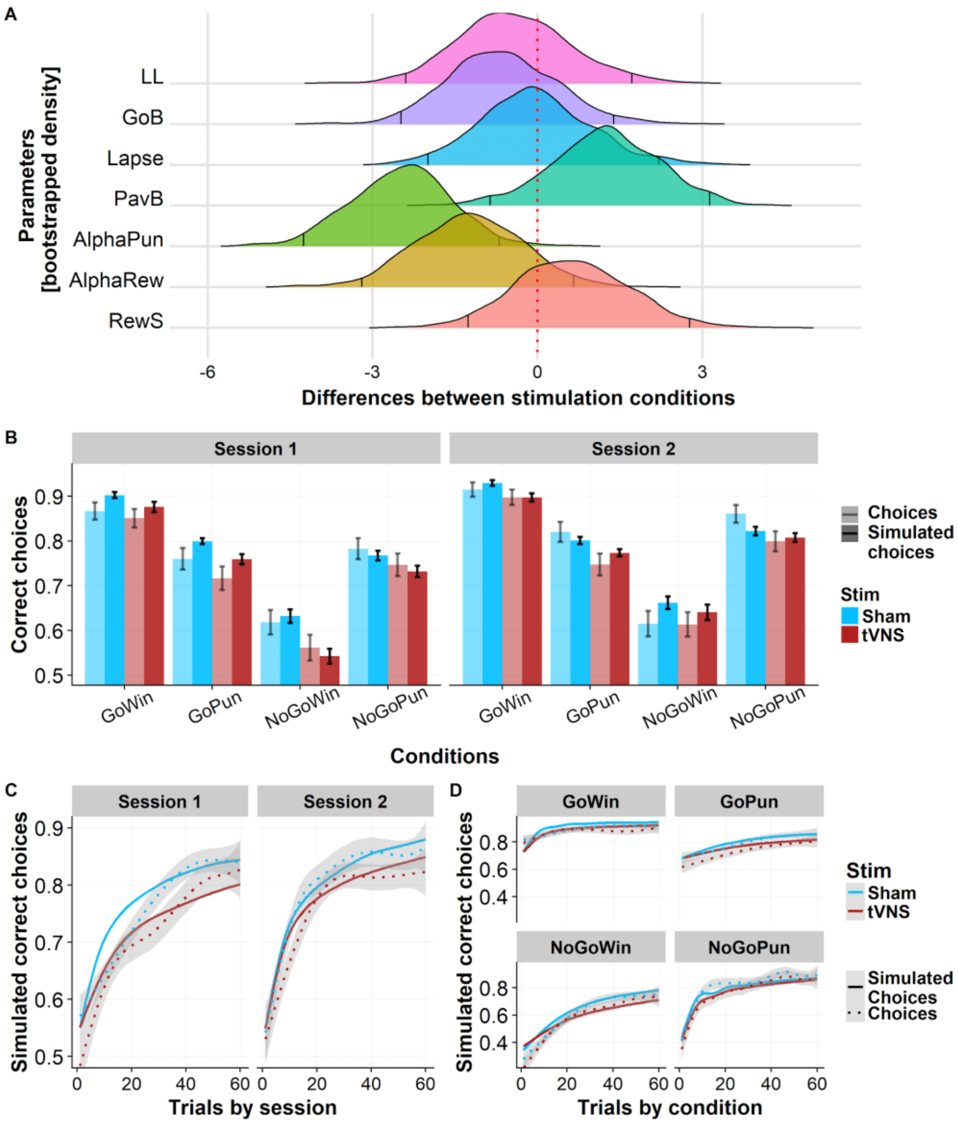


**Figure S.1:** Reductions in learning rate are driven by slowed learning in punishment conditions. A: Bootstrapped density plots of differences in individual parameter estimates between tVNS and sham stimulation. Lines indicate 95% confidence intervals. B: Mean choice accuracy simulated from individual parameter estimates recover stimulation effects in each session and condition. Error bars depict 95% confidence intervals. Transparent bars represent empirical data, dark bars simulated data C: Choices simulated from individual parameter estimates recover participants' choice patterns and stimulation effects for sessions. D: Recovered choice patterns indicate no difference in stimulation effects depending on valence or action. Fit between empirical and simulated data is comparable between the six- and five parameter model (B-D; Figure 3 C-E). LL = Log-Likelihood, GoB = Go bias, PavB = Pavlovian bias (π), RewS = Reward sensitivity (ρ)

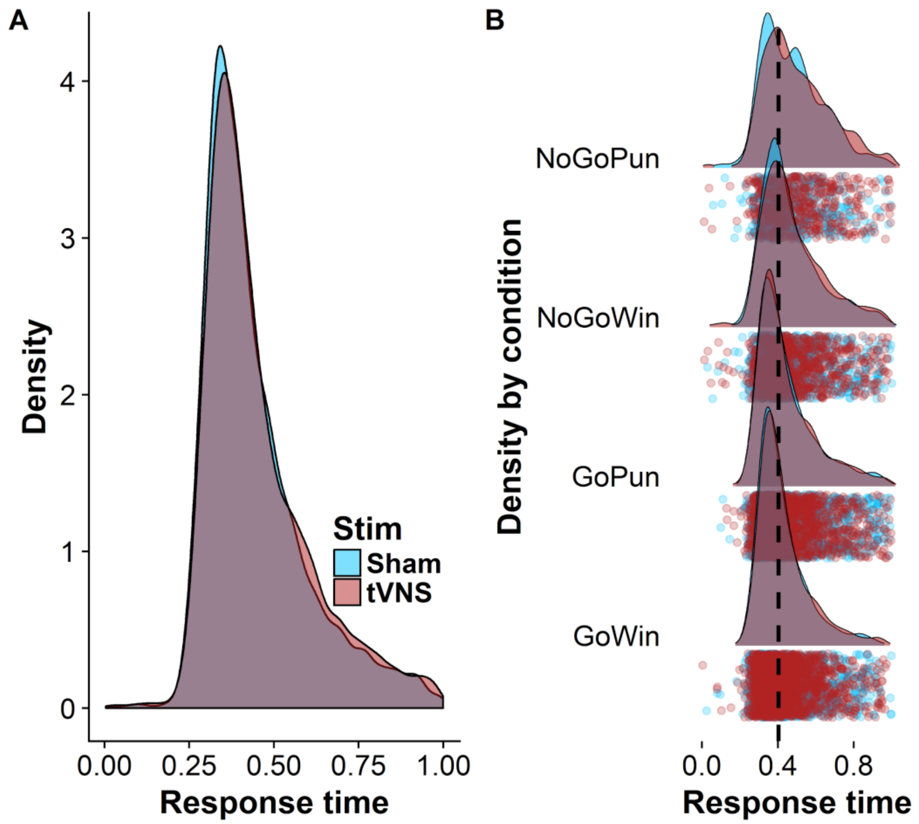


**Figure S.2.** Transcutaneous vagus nerve stimulation (tVNS) does not induce changes in response time as is seen across (A) and within (B) conditions.
